# Comparative genomics-first approach to understand diversification of secondary metabolite biosynthetic pathways in symbiotic dinoflagellates

**DOI:** 10.1101/376251

**Authors:** Girish Beedessee, Kanako Hisata, Michael C. Roy, Frances M. Van Dolah, Noriyuki Satoh, Eiichi Shoguchi

## Abstract

Symbiotic dinoflagellates of the genus *Symbiodinium* are photosynthetic and unicellular. They possess smaller nuclear genomes than other dinoflagellates and produce structurally specialized, biologically active, secondary metabolites. Polyketide biosynthetic genes of toxic dinoflagellates have been studied extensively using transcriptomic analyses; however, a comparative genomic approach to understand secondary metabolism has been hampered by their large genome sizes. Here, we use a combined genomic and metabolomics approach to investigate the structure and diversification of secondary metabolite genes to understand how chemical diversity arises in three decoded *Symbiodinium* genomes (A3, B1 and C). Our analyses identify 71 polyketide synthase and 41 non-ribosomal peptide synthetase genes from two newly decoded genomes of clades A3 and C. Additionally, phylogenetic analyses indicate that almost all of the gene families are derived from lineage-specific gene duplications in *Symbiodinium* clades, suggesting divergence for environmental adaptation. Few metabolic pathways are conserved among the three clades and we detect metabolic similarity only in the recently diverged clades, B1 and C. We establish that secondary metabolism protein architecture guides substrate specificity and that gene duplication and domain shuffling have resulted in diversification of secondary metabolism genes.

## Introduction

Dinoflagellates of the genus *Symbiodinium* exist freely in the water column and in symbiotic associations with many invertebrates, such as corals, clams, and anemones. This invertebrate-Symbiodinium mutualism seems to provide a competitive advantage [1], resulting in the production and exchange of metabolites by both organisms [2]. Members of this genus are sources of unusual, large, polyhydroxyl and polyether compounds or so-called “super-carbon-chain compounds (SCC),” composed of long-chain backbones functionalized by oxygen [3]. It is possible to classify diverse *Symbiodinium* into nine clades (A to I) by molecular phylogenetic analysis [4]. Several of these compounds such as zooxanthellatoxins (ZTs) and zooxanthellamides (ZADs) have been isolated from several *Symbiodinium* clades and a clade-to-metabolite relationship has been proposed and experimentally supported, in which strains of specific *Symbiodinium* clades produce specific metabolites [5]. Nakamura *et al*. [6] suggested the existence of shared biogenetic processes, such as the polyketide pathway with glycine as the starting substrate, yielding products with structural similarities to palytoxins and zooxanthellatoxins. Over the years, other secondary metabolites have been isolated from these clades, but their ecological roles and biosynthetic pathways have yet to be identified [7]. A preliminary genomic survey reported the presence and organization of secondary metabolite genes in *Symbiodinium minutum*, overcoming limitations of previous transcriptomic surveys [8]. Availability of new *Symbiodinium* genomes now allows us to survey and compare genes associated with metabolite biosynthesis [9–12]. However, how chemical diversity arises within *Symbiodinium* clades is still unknown. Evolution of novel chemistry depends on diversity-generating metabolism, which comprises broad-substrate enzymes [13]. Metabolic pathways accept many different substrates, generating diverse chemical products and this provides organisms with unique chemistry to face environmental challenges [14]. Polyketide synthase (PKS) and non-ribosomal peptide synthase (NRPS) are two important classes of such modular enzymes involved in secondary metabolite biosynthesis, where modules integrate building blocks into a growing chain like an assembly line [15].

PKSs comprise three core domains: an acyl-transferase (AT) domain that recognizes and loads small carboxylic acid building blocks, an acyl-carrier protein (ACP) domain that retains the building blocks, and a ketosynthase (KS) domain that builds the polyketide chain via condensation with optional domains such as ketoreductase (KR), dehydratase (DH), and enoylreductase (ER) domains [16]. Polyketide synthases are also closely related to fatty acid synthases (FASs) and share the same core of enzymatic activities, implying a common evolutionary history [17]. On the other hand, NRPSs are modular multi-enzyme complexes that synthesize a diverse array of biological active peptides or lipopeptides [18]. Biosynthesis of non-ribosomal peptides occurs via the action of catalytic modules within NRPS that are composed of three compulsory domains: adenylation (A), thiolation (T), and condensation (C). The process involves recognition of an amino acid (or hydroxyl acid) by the A-domain, covalent attachment of the adenylated amino acid to a phosphopantetheine carrier of the T-domain, and finally peptide bond formation between two consecutively bound amino acids to a growing peptide chain by the C-domain. These core domains are often supported by domains such as an epimerization (E) domain, a dual/epimerization (E/C) domain, a reductase (R) domain, a methylation (MT) domain, a cyclization (Cyc) domain or an oxidation (Ox) domain [19]. Finally, PKSs and NRPSs have a common thioesterase (TE) domain that releases assembled polyketide and polypeptide chains, respectively, from the enzyme complex. PKS and NRPS pathways often cross-talk such that a polyketide product is elongated by NRPS or *vice versa* to produce hybrid natural products, thereby increasing structural diversity [20]. Based on protein organization, PKSs are further categorized into three (Type I, II and III) types, and FASs into two (Type I and II) [21].

Pathways involved in secondary metabolite biosynthesis are among the most rapidly evolving genetic elements [22]. Processes such as gene loss, duplication, and horizontal gene transfer (HGT) have played significant roles in distribution of PKSs in fungi and bacteria [23, 17]. Mutations, domain rearrangements, and module duplications within PKS and NRPS genes account for generation of novel, diverse small-molecules [22]. There exist several entry points where combinatorial potential can arise. In PKS, the AT domain shows specificity for malonyl-CoA, methylmalonyl-CoA, or other malonyl-CoAs, while the KR domain can create two stereoisomers [24]. In contrast, NRPS can accept 500 different monomers, including nonproteinogenic amino acids, fatty acids and α-hydroxyl acids [25, 26]. Different tailoring enzymes, like glycosyltransferases, halogenases, methyltransferases, and oxidoreductases can further alter the chemical structure of secondary metabolites by addition of various functional groups [27].

To investigate existence of shared biosynthetic pathways, we cultured three *Symbiodinium* strains (clades A3, B1, and C) known to produce different metabolites, and we surveyed their genomes [9, 12] for genes involved in polyketide and non-ribosomal peptide biosynthesis. Additionally, we examined how these genomes are equipped to expand their gene repertoire for biosynthesis of complex secondary metabolites and suggest possible diversification mechanisms that may contribute to such chemical variability and modularity.

## Materials and Methods

### *Symbiodinium* cultures

Clade B1 was isolated from the stony coral, *Montastraea (Orbicella) faveolata* by Dr. Mary Alice Coffroth (University of New York, Buffalo, USA) and clades A3 and C were isolated from the clam *Tridacna crocea* and bivalve *Fragum* sp., respectively, by Late Dr. Terufumi Yamasu (University of the Ryukus, Okinawa, Japan). Cultures were maintained in autoclaved, artificial seawater containing 1X Guillard’s (F/2) marine-water enrichment solution (Sigma-Aldrich: GO 154), supplemented with antibiotics (ampicillin (100 μg/mL), kanamycin (50 μg/mL), and streptomycin (50 μg/mL)). Culturing and sampling were performed according to the protocol of Shoguchi *et al*. [9].

### Data retrieval

The genome browser MarinegenomicsDB (http://marinegenomics.oist.jp/genomes/gallery/) and the *Symbiodinium kawagutii* browser (http://web.malab.cn/symka_new/genome.jsp) were accessed in order to retrieve PKS (KS & AT), FAS (FabB-KASI, FabF-KASII & FabD) and NRPS (A & C) sequences from clades A3, B1, and C and *Symbiodinium kawagutii*, respectively [10,12,28]. In addition, transcriptome datasets for several dinoflagellates, apicomplexans, stramenopiles, and haptophytes were downloaded from the Marine Microbial Eukaryote Transcriptome Sequencing Project (MMETSP) (http://datacommons.cyverse.org/browse/iplant/home/shared/imicrobe/camera) and surveyed for comparative analysis [29]. Amino acid sequences of several other prokaryotes, fungal, animal, and chlorophyte PKS and NRPS domains were obtained from NCBI Genbank with additional sequences from dinoflagellates [30, 31]. Further NRPS sequences from *Proteobacteria*, *Firmicutes*, and *Cyanobacteria* were obtained from Wang *et al*. [15]. Functional prediction and conserved active site residues in sequences were identified using Pfam [32]. Only PKS, FAS, and NRPS sequences with full domains and conserved active sites were used in the analysis. Throughout this manuscript, we name gene models from the three *Symbiodinium* clades (A3, B1 and C) with the letters A, B, and C so as to improve the readability and interpretation. Details of gene IDs and their transcriptome support are provided in Supplementary Tables S1-S4.

### Phylogenetic analysis

Type I and II PKS/FAS and condensation (C) & adenylation (A) domain sequences representing different taxa were used for Bayesian inference and maximum likelihood analysis. Four amino acid (aa) domain sequence datasets comprising of 233 KS sequences (226 aa), 96 AT sequences (208 aa), 117 A-sequences (400 aa), and 110 C-sequences (260 aa) were aligned using the MUSCLE algorithm [33]. Sites within alignments where homology was ambiguous (e.g. large insertions and deletions) were removed prior to phylogenetic analyses. Maximum likelihood phylogenetic analysis was performed using RaxML with 1000 bootstraps using the GAMMA and Le-Gasquel amino acid replacement matrix [34]. Bayesian inference was conducted with MrBayes v.3.2 [35] using the same replacement model and run to maximum of six million generations and four chains or until the posterior probability approached 0.01. Statistics and trees were summarized using a burn-in of 25% of the data. Using two methods provided a convenient way to verify different phylogenetic estimates, since each method has its intrinsic strengths and assumptions about the evolutionary process. Trees were edited using Figtree (http://tree.bio.ed.ac.uk/software/figtree/).

### *In silico* analysis of PKS and NRPS genes and genomic locations

Monomer prediction based on specificity of the A-domain was determined using the Latent Semantic Indexing of the LSI-based A-domain predictor [36]. NaPDos was used to determine C-domain types [37]. For AT domains, *Symbiodinium* sequences were compared to the Hidden Markov Model-based ensemble (HMM) generated by Khayatt *et al*. [38]. Additional information about substrate specificity was detected using I-TASSER [39]. AntiSMASH (Antibiotics & Secondary Metabolite Analysis SHell) version 4.1.0 was used with default settings to identify NRPS and PKS gene clusters within scaffold regions using nucleotides sequences as queries [40]. Subcellular localization of PKS gene products toward organelles (e.g. chloroplast and mitochondria) or the presence of signal peptide or membrane anchor was determined using ChloroP 1.1 and TargetP 1.1 using a cut-off score of ≥ 0.50 each and the subcellular localization predictor, DeepLoc [41–43]. NUCmer operation of SyMap v4.2 (Synteny Mapping and Analysis Program) was used to align and visualize syntenic relationships between the three *Symbiodinium*clades [44]. Scaffold information and descriptions of these genomes were imported into SyMap as GFFs (General Feature Files). To determine orthologs, we performed an all-against-all BLAST search of *PKS*-coding scaffolds of one genome against itself at a BLAST bit score cutoff of ≥ 100 and e-value ≤ e^-20^. Outputs were parsed and processed, and orthologous pair detection was conducted using custom perl scripts. Possible segmental duplications were visualized using Circos [45]. GC content variations in PKS-coding scaffolds were analyzed using GC-profile with a halting parameter of 100 [46]. LTR Finder 1.05 was used with defaults parameters to search for long terminal repeat (LTR) retrotransposon-specific features [47].

### Polyol extraction from *Symbiodinium* cultures (A3, B1, and C)

All cultured biomass samples (A3, B1, and C) were treated as previously described [8]. Cultured cells were collected by centrifugation (9,000 × g and 14,000 g, 10 min, 10 °C). After discarding the supernatant, a cell pellet was extracted with methanol (three times) at room temperature. Methanol (400 μL) was added to the biomass followed by vortexing (1 min), sonication (10 min), and centrifugation (14,000 × g, 10 min, 10 °C) to give a methanol extract. The resulting clear solution was transferred into a new tube. By adding methanol (400 μL) to the residue, a 2^nd^ methanol extraction was conducted in the same fashion. The 2^nd^ clear methanol extract was again collected and stored at -30 °C. Additional methanol (400 μL) was added to the residue, vortexed (1 min), and kept overnight at room temperature. After centrifugation, the 3^rd^ methanol extract was pooled with the previous extracts (total 1,200 μL), and marked as crude extract. To remove lipophilic materials from the crude extract, an aliquot (50 μL) of the crude extract was suspended in 50 μL water-methanol (90:10) containing 0.5% formic acid. The suspension was vortexed (30 sec) and centrifuged (14,000 × g, 10 min, 10 °C) to give a clean solution. The clean solution was transferred into a new tube (the stock solution) and the insoluble part was discarded. The stock solution was kept at -30°C before NanoLC-MS analysis or immediately analyzed after dilution. All crude extracts were lyophilized and stored at -30 °C.

### NanoLC-MS analysis of the *Symbiodinium* methanol extract

A Thermo Scientific hybrid (LTQ Orbitrap) mass spectrometer was used for MS data collection. The mass spectrometer was equipped with a HPLC (Paradigm MS4, Michrom Bioresources Inc.), an auto-sampler (HTC PAL, CTC Analytics), and a nanoelectrospray ion source (NSI). The high-resolution MS spectrum was acquired at 60,000 resolution in FTMS mode (Orbitrap), full mass range *m/z* 400-2,000 Da with capillary temperature (200 °C), spray voltage (1.9 kV), and both positive and negative ion modes were used. The lipid-depleted crude extract (stock solution) was diluted (1:50) by adding water-methanol (50:50) containing 0.25% formic acid and separated on a capillary ODS column (50 × 0.18 mm, 3 μm, C_18_, Supelco). A 20-min gradient (10% B for 0-2 min, 10-100% B for 2-10 min, hold 100% B for 10-15 min, equilibration 10% B for 15.1-20.0 min, where solvent A was water: acetonitrile 98:2 and solvent B was water: acetonitrile 2:98, both containing 0.1% formic acid; flow rate 2.0 μL/min, injection, 2.0 μL) was used for polyol separation.

### KS protein localization

KS protein localizations were visualized using a modified version of the protocol of Berdieva *et al*. [48]. Briefly, *Symbiodinium* cells were prefixed in methanol: F/2 medium (1:1) at RT for 15 min. Samples were then fixed in methanol at -20 °C overnight. Cells were washed in PBS, followed by permeabilization with 1% Triton X-100 for 15 min (5 min for clade B1), further washed with PBS and blocked with 5% normal goat serum-PBST for 1h. Subsequently cells were incubated overnight at 4 °C with primary anti-KS antibodies at a 1:100 dilution in blocking solution. Primary antibody solution was then removed with 3 × 5-min PBS washes and cells were incubated with Alexa Fluor 488 (Abcam Cat #ab150077) secondary antibody for 1h at RT (1:100 in blocking solution), ending with several PBS washes. Coverslips were mounted in DAPI-containing Vectashield on glass slides and visualized using a Zeiss Axio-Observer Z1 LSM780 confocal microscope under a Plan-APOCHROMAT 63X/1.4 oil DIC objective lens. Fluorescence excitation/emission wavelengths were 410/482 nm for DAPI, 499/614 nm for Alexa Fluor 488, and 649/740 nm for chlorophyll autofluorescence. Data were acquired using Zeiss ZEN version 14.0.8.201 software. For negative controls, primary antibodies were omitted. Z-stacks profiles were analyzed using ImageJ [49]. DIC imaging was performed using a Zeiss Image-Z1 under 40X.

## Results

### Phylogenetic and syntenic analyses of ketosynthase and acyltransferase domains

To understand diversification and molecular evolution of PKS and FAS, we performed an extensive search for PKS (KS & AT) and FAS (FabB-KASI, FabF-KASII & FabD) genes within three *Symbiodinium* genomes, as these domains are conserved [23]. We integrated KS, FabB-KASI, and FabF-KASII domains into a dataset of well-characterized sequences from several taxa and subjected these data to phylogenetic analysis. The tree shows that the majority of KS domains clustered according to their domain organization types under a reliable node (Fig. 1, Supplementary Figure S1-S2). Recently, Kohli *et al*. [31] described contigs encoding multiple PKS domains in the dinoflagellates, *Gambierdiscus excentricus* and *Gambierdiscus polynesiensis*. Those sequences were also included in the dataset and clustered into Dinoflagellate PKS II clade (blue highlighted inset of Fig. 1). From this survey, we confirmed the presence of 25 KS sequences each from clades A3 and C. Our analysis showed only one gene model (B1030341.t1) associated with type II fatty acid synthesis (FabF-KASII) and one gene model (B1027279.t1) in the FabB-KASI clade. In general, our analysis mirrors that of Kohli *et al*. [50], where there is a demarcation between type II FAS and Type I PKS & FAS. In addition, our analysis reveals the expanded nature of KS genes into nine PKS clades (Dinoflagellate PKS I-III and *Symbiodinium* PKS I-V) associated with either mono- or multifunctional domains. Interestingly, one clade (Dinoflagellate PKS-I) was closely related to cyanobacterial KS sequences. Scanning the GC profile of PKS-I clade scaffolds of clade C showed some regions of higher GC content (45-46.5%), compared to the average genomic GC content of 43.0%, indicative of gene transfer (Supplementary Figure S3). ~3% (3/83) of the sequences contain the cTP (chloroplast transit peptide) signal while 12% (10/83) contained mitochondrial targeting peptide (mTP) or secretory signal each (Fig. 1, Supplementary Table S5).

**Fig. 1.**
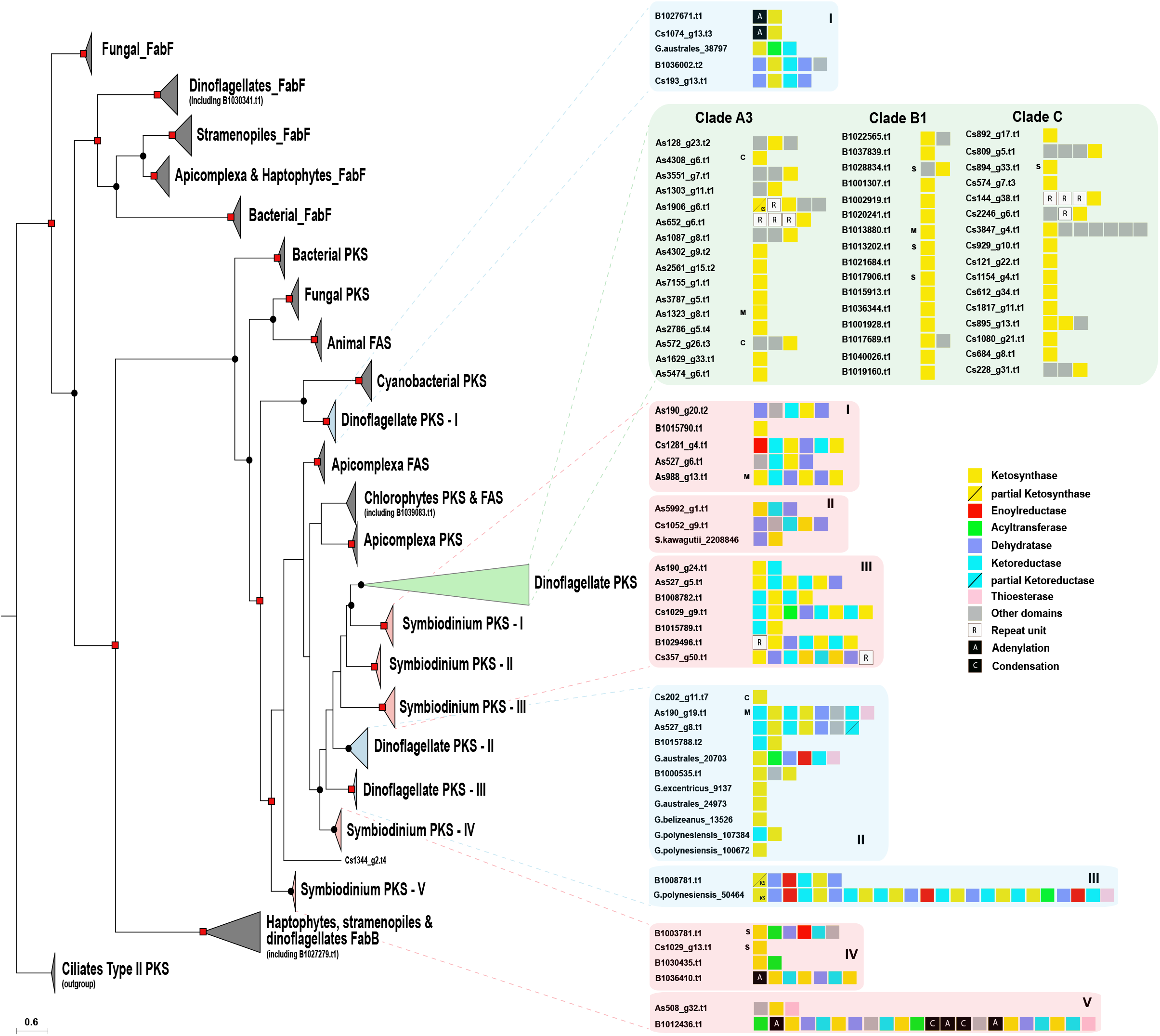
Phylogenetic tree of ketosynthase (KS) domains of prokaryotic and eukaryotic polyketide and fatty acid synthases. Analysis of ketosynthase, FabB-KASI, and FabF-KASII domains show diversification of these domains into nine clades, comprised of mono-and multifunctional domains. Dots and squares indicate posterior probabilities of 0.70-0.89 and 0.9-1.0, respectively, generated by Bayesian inference. Inserts provide details of sub-clades as well as gene model architecture. C, M, and S denote chloroplasts, mitochondria, and secretory signal peptides, respectively.

A striking feature among the three genomes is the high number (26) of *trans*-AT genes in contrast to *cis*-AT (4). A phylogenetic tree of the AT domain consisted of two main *Symbiodinium* clades, *cis*-AT and *trans*-AT (Fig. 2, Supplementary Figure S4-S5), deviating from the classical substrate-based clustering [38]. Alignment of the *trans*-AT motif revealed a deviation from the usual GHSxG conserved motif to GLSxG where x can be any residue; thus a change from a basic amino acid (histidine) to an aliphatic one (leucine) while *cis*-AT maintained their GHSxG motif. The implication of His➔Leu remains to be investigated (Fig. 2). Use of the HMMs by Khayatt *et al*. [38] didn’t suggest any clear distinction regarding which substrates are being incorporated into biosynthetic pathways; however, I-TASSER predicted that most *Symbiodinium* AT sequences pertain to the family of malonyl-CoA ACP transferase (data not shown). Downstream of the active site serine, a motif (YASH or HAFH) is involved in the choice of either methylmalonyl-CoA or malonyl-CoA, respectively [51]. The motif, GAFH, present in most *Symbiodinium* sequences reflects the prediction of I-TASSER. ~9 % (3/33) of AT gene models contained the cTP or mTP signals (Fig. 2).

**Fig. 2.**
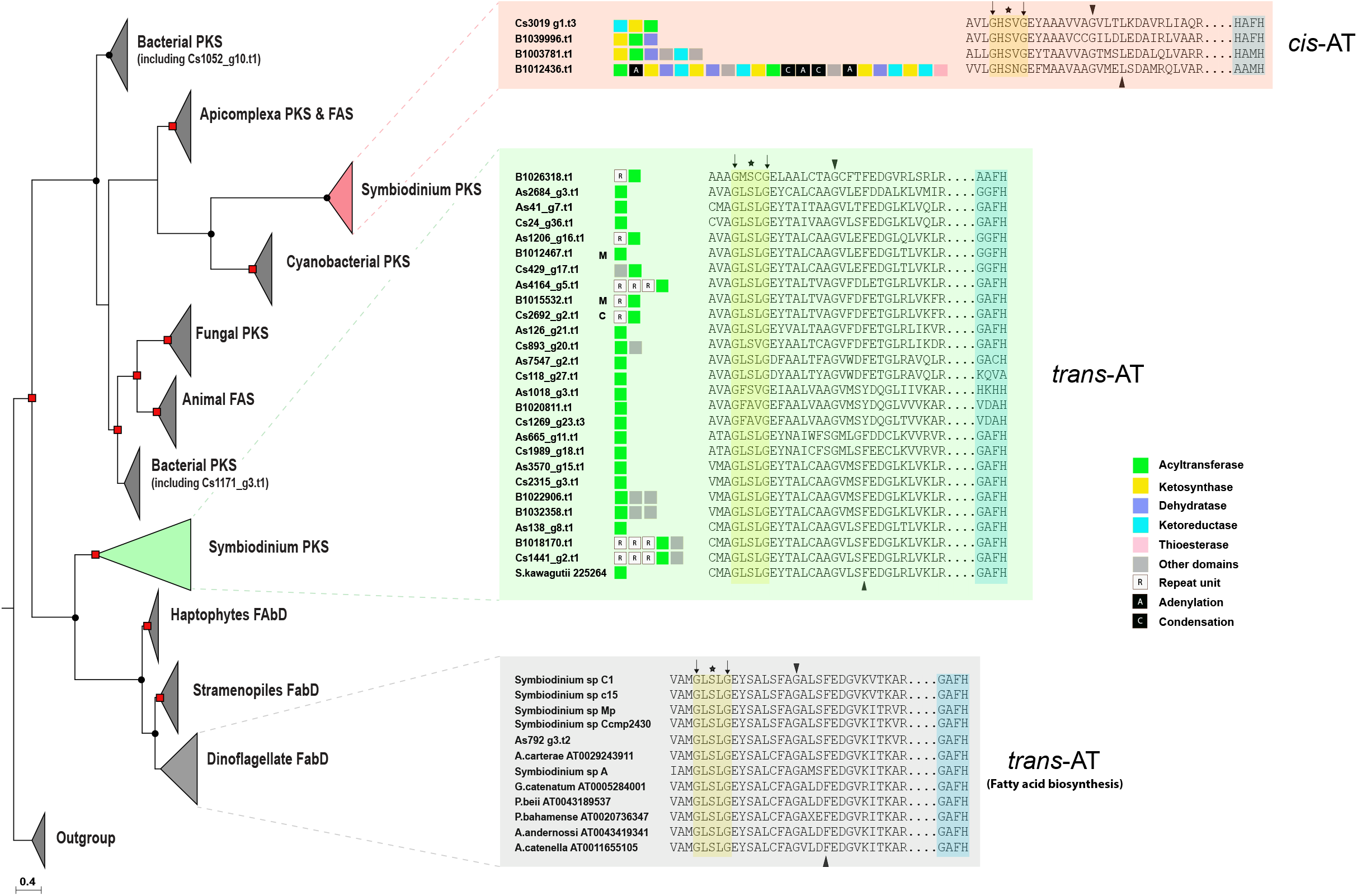
Phylogenetic tree of acyltransferase (AT) domain of prokaryotic and eukaryotic polyketide and fatty acid synthases. Analysis of acyltransferase domain show clear demarcation between *cis*- and *trans*-AT. Dots and square indicate posterior probability 0.70-0.89 and 0.9-1.0, respectively generated by Bayesian inference. Details of sequences are provided in box inserts. Asterisk indicates active site residue, black triangles indicate conserved residues characteristic for specific substrate groups, and black arrows indicate overall conserved residues used by HMM (Khayatt *et al.*, 2013). C, M and S depicts chloroplast, mitochondria, and secretory signal peptide respectively.

Comparative visualization of *PKS*-containing scaffolds from the three genomes showed extensive duplication events in the three clades between genes associated with polyketide biosynthetic clusters (Fig. 3). Genomic synteny was observed between clades B1 and A3 (8 syntenic blocks), clades B1 and C (10 syntenic blocks) and clades A3 and C (7 syntenic blocks) (Fig. 3b-d) while only four *PKS*-containing gene clusters were found to be shared among all three clades (green boxes in Fig. 3b-d). The observed rearrangements within the syntenic scaffolds included mainly deletions. Transposons were found on scaffolds carrying *PKS*- and *NRPS*-encoding genes, suggesting that these genes can be influenced by transposable elements. 47% (52/110) of *PKS*- and 34% (14/41) *NRPS*-containing scaffolds possessed LTR signatures (Supplementary Table S6-S7). Taken together, these results indicate that PKS genes have diversified in each *Symbiodinium* clade by several evolutionary processes.

**Fig. 3.**
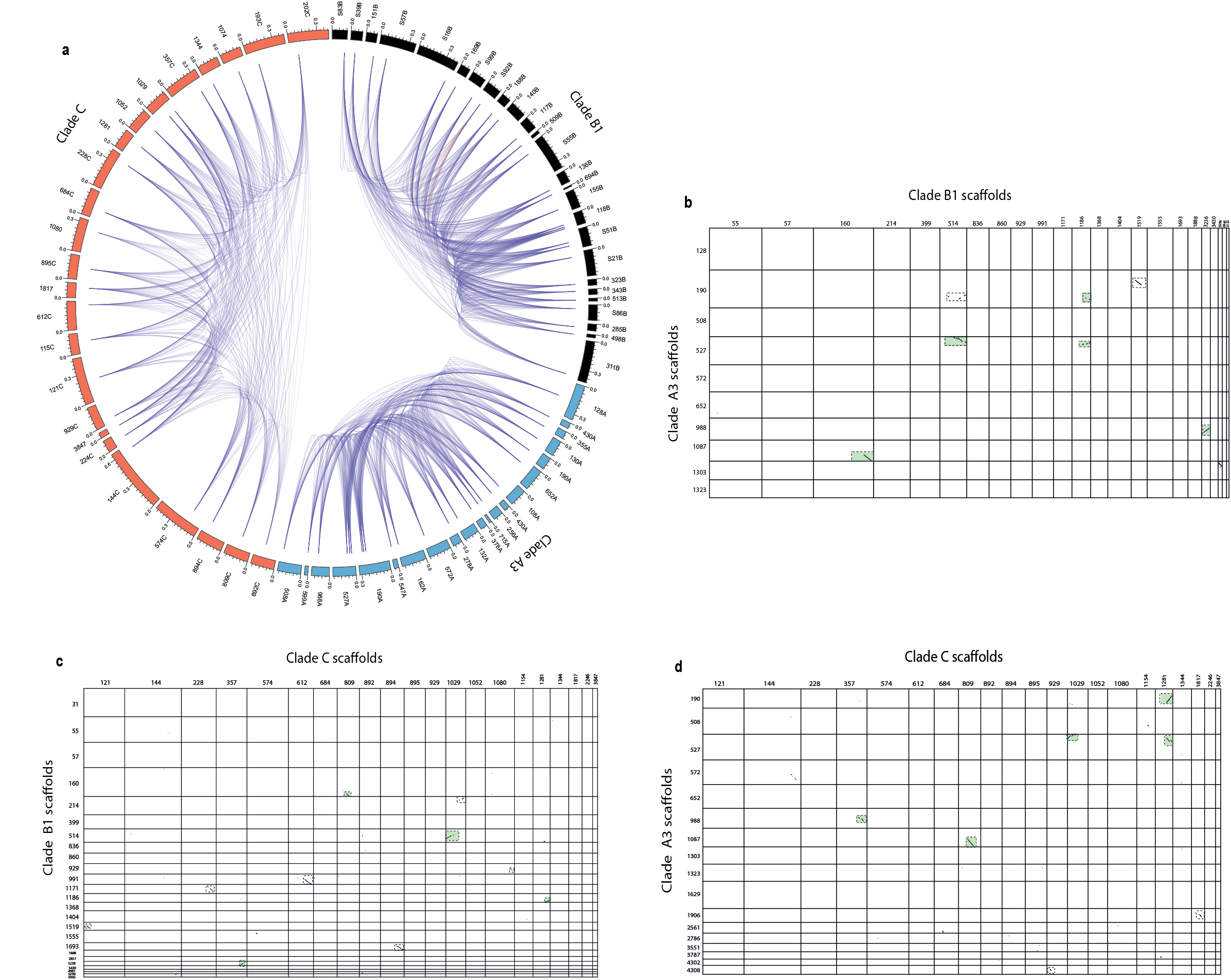
Pathway duplication and conservation within and across *Symbiodinium* clades. **a** Plot showing duplicate gene distribution within PKS-containing scaffolds of three *Symbiodinium*genomes. Colored sections (black = clade B1, orange = clade C, blue = clade A3) represent scaffolds studied in Fig. 1. A link represents a possible duplication event between two domains. **b** Synteny plot of clade A3 and B1 PKS-containing scaffolds. **c** Synteny plot of clade B1 and C PKS-containing scaffolds. **d** Synteny plot of clade A3 and C PKS-containing scaffolds. Dotted boxes highlight regions of significant homology between genomes. Green colored dotted boxes show common regions shared among the three genomes.

### Phylogenetic analysis of adenylation and condensation domain subtypes (^L^C_L_, ^D^C_L_, Cyc and dual E/C) in NRPS proteins

To understand if freestanding A-domains identified in *Symbiodinium* genomes follow the same non-ribosomal code of traditional NRPS systems [52], we performed a phylogenetic comparison involving 117 adenylation sequences from different taxa. One major observation was that a freestanding A-domain from *Symbiodinium* appears in three major clades that utilize tryptophan, glycine, and phenylalanine as substrates (three highlighted clades in Fig. 4a, Supplementary Figure S6-S7). In contrast, other proteins with di- or multi-domains displayed affinity for various substrates. Phylogenetic analysis of condensation domains was dominated by functional categories of C-domains rather than species phylogeny or substrate specificity alone. Four specific functional categories were clearly supported namely (1) ordinary C-domains that are composed of ^L^C_L_ and ^D^C_L_, (2) heterocyclization (Cyc) domains, (3) dual E/C domains and (4) starter domains, which are found on initiation modules (Fig. 4b, Supplementary Figures S8 and S9). It is clear that *Symbiodinium* clades are rich in ^L^C_L_ subtypes, that catalyze the condensation of two L-amino acids, in contrast to a ^D^C_L_ that links an L-amino acid to a D-amino acid. Both catalysts possess a conserved His-motif in their active sites with a consensus sequence of HHxxxDG, where x can be any residue. Our survey revealed the presence of six condensation domains with the consensus motif being maintained, except for G being substituted with L and N in B1036245 .t1 and Cs535_g6.t1, respectively (data not shown). The phylogeny also supports the close relationship between ^L^C_L_ and starter C domains and that of dual E/C and ^D^C_L_ domains, as previously reported in bacterial genomes, confirming the reliability of our analysis [53]. These results demonstrate the specificity of NRPS genes for specific amino acids, thus introducing a degree of chemical diversity in non-ribosomal peptide biosynthesis.

**Fig. 4.**
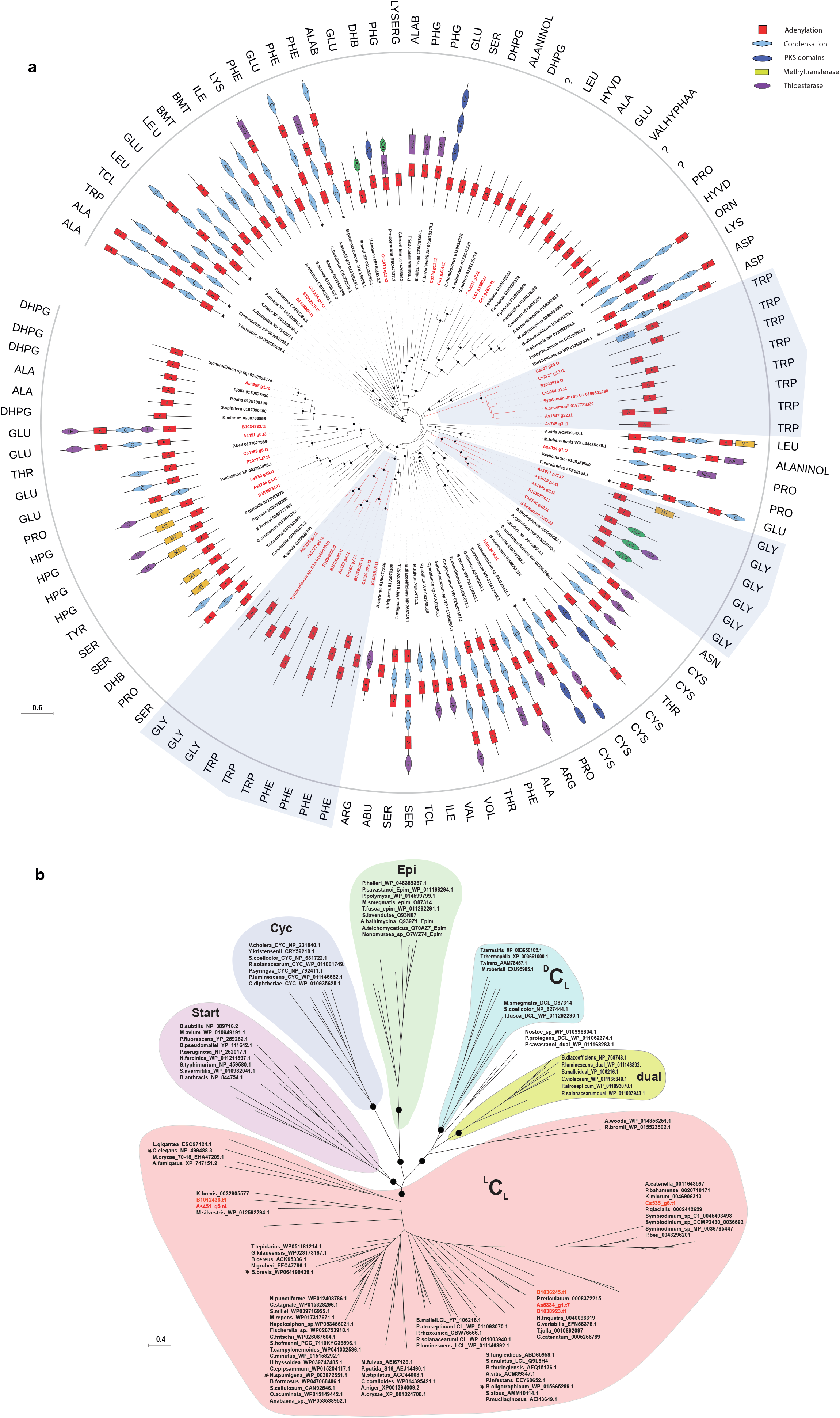
Phylogenetic analysis of adenylation (A) and condensation (C) domains of prokaryotic and eukaryotic NRPS. Dots indicate posterior probability ≥ 0.70 generated by Bayesian inference. **a** Analysis of adenylation domains show specificity of monofunctional domains from *Symbiodinium* toward three amino acids (glycine, tryptophan, and phenylalanine) as indicated by shaded regions. Asterisks indicate multifunctional proteins that are too long to display. Details of protein sequences are provided in Supplementary Table S4. **b** Condensation domains from *Symbiodinium* belong to the ^L^CL type (shown in red). Asterisks indicate sequences with different specificities beside clade subtype specificity.

### Identification of metabolites and biosynthetic gene clusters from *Symbiodinium* genomes

Polyols were identified based on their high-resolution mass data, as summarized in Beedessee *et al*. [8]. Doubly charged ions (negative ions) were searched for larger polyols (>2600 Da) in the MS spectra. Sample A3 showed the presence of zooxanthellatoxin-B (ZT-B), albeit in small amount, with an m/z of 1414.74 for the [M-2H]^2-^ (Supplementary Figure 10a). Only zooxanthellamide D (ZAD-D) could be identified from sample B1 with extracted ions at *m/z* 1050.57 for the [M+H]^+^ (Supplementary Figure 10b). No SCCs could be identified from sample C despite presence of many polyols (Supplementary Figure 10c-11a). Samples B1 and C also showed similar LC-MS profiles and contained some identical unknown SCCs in the molecular weight range of 2,600-2,850 Da (Supplementary Figure 11b). It should be noted that other polyhydroxy SCCs were also detected in the crude methanol extracts of all samples and none of them corresponds to known zooxanthella polyhydroxy molecules [8]. Purification and characterization of these unknown polyhydroxy SCC compounds could be interesting. Analysis using antiSMASH on *Symbiodinium* genomes matched four PKS-NRPS -containing clusters to known biosynthetic gene clusters, with similarities between 25-46% (Fig. 5a). Clade A3 harbors a gene cluster with similarity to ajudazol and phenalamide biosynthetic genes from *Streptomyces* species and *Chondromyces crocatus* while clade B1 shares similarity with a phenalamide biosynthetic cluster from *Chondromyces crocatus*. High sequence similarity was noted in clade A3, offering an example of module duplication between modules of gene models in one scaffold, as well as between modules of different scaffolds (Fig. 5b). In order to examine the localization of KS protein, antibodies against the KS domain were used. Immunolocalization showed that KS proteins were mainly associated with reticulate chloroplasts in *Symbiodinium* clade C (Supplementary Fig. 12), although the possibility remains that KS proteins are localized in other organelles. Similar observations on the location of KS proteins in chloroplasts have been reported in *Karenia brevis* [54].

**Fig. 5.**
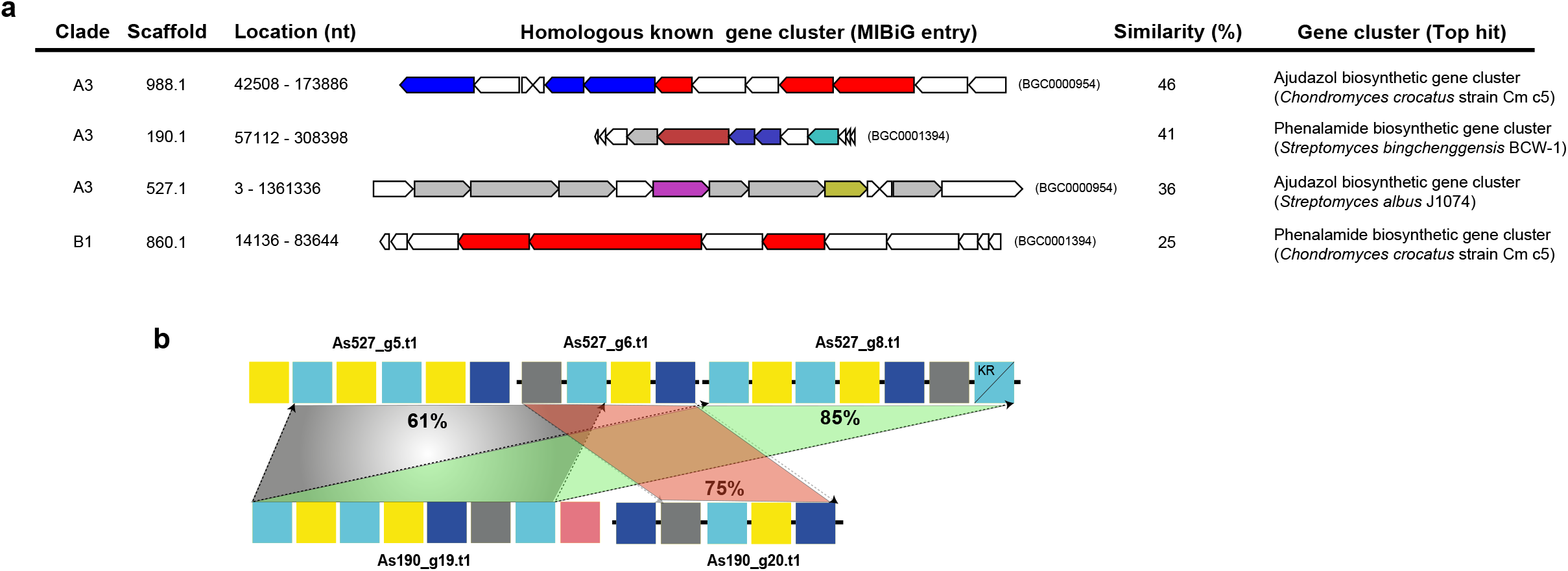
Multifunctional PKS genes in *Symbiodinium*. **a** Table showing homologous gene clusters and similarities of different scaffolds from *Symbiodinium* clades obtained using antiSMASH version 4.1.0. Details of each gene cluster can be obtained using the MIBiG (Minimum Information about a Biosynthetic Gene cluster) entry number and is accessible at https://mibig.secondarymetabolites.org/repository.html **b** Homology comparison of two scaffolds (527.1 and 190.1 of Clade A3) shows an example of module duplication. Numbers indicate the percentage of identity shared between sequences. Details of modules are depicted in Fig. 1.

## Discussion

### Evolution of modularity within *Symbiodinium* genomes

The genomic analysis at the clade-level presented here reveals expanded genetic diversity of metabolite-producing capacity in *Symbiodinium* dinoflagellates. The polyketide biosynthesis machinery gains its functional and genetic modularity by changes through combinatorial events assisted by gene duplication, horizontal gene transfer (HGT), and recombination [55]. The presence of a large number of monofunctional KS or AT domains within these genomes raises questions about the evolution of modularity. Our analysis shows that domain as well as module duplications established an important evolutionary mechanism toward modularity (Fig. 5b). Dinoflagellate genomes are scattered with large numbers of repeats, with frequent recombination events, and possess genes with high copy numbers due to duplication [9–11]. These features might have led to decomposition of Type I multifunctional PKS clusters, a phenomenon involving shuffling of domains and modules previously observed in bacteria [17]. However, there is increasing evidence of multifunctional PKS domains in several dinoflagellates, indicating that multifunctionality coevolves with monofunctional domains [8, 31, 56]. Our data show that monofunctional PKSs are related to multifunctional PKS (Fig. 1) but it remains unclear whether fusion of monofunctional PKS domains led to multifunctionality or *vice versa*. Retrotransposons may have been important contributors in the expansion of PKS and NRPS, since 34% and 47% of the scaffolds, respectively, are predicted to contain LTR signatures (Fig. 6, Supplementary Table S6). Retrogenes account for >20% of all genes in *Symbiodinium* clades [57]. The Ty1/copia LTR retrotransposon has been proposed as a likely candidate driver for retroposition in *Oxyrrhis marina* [58].

**Fig 6.**
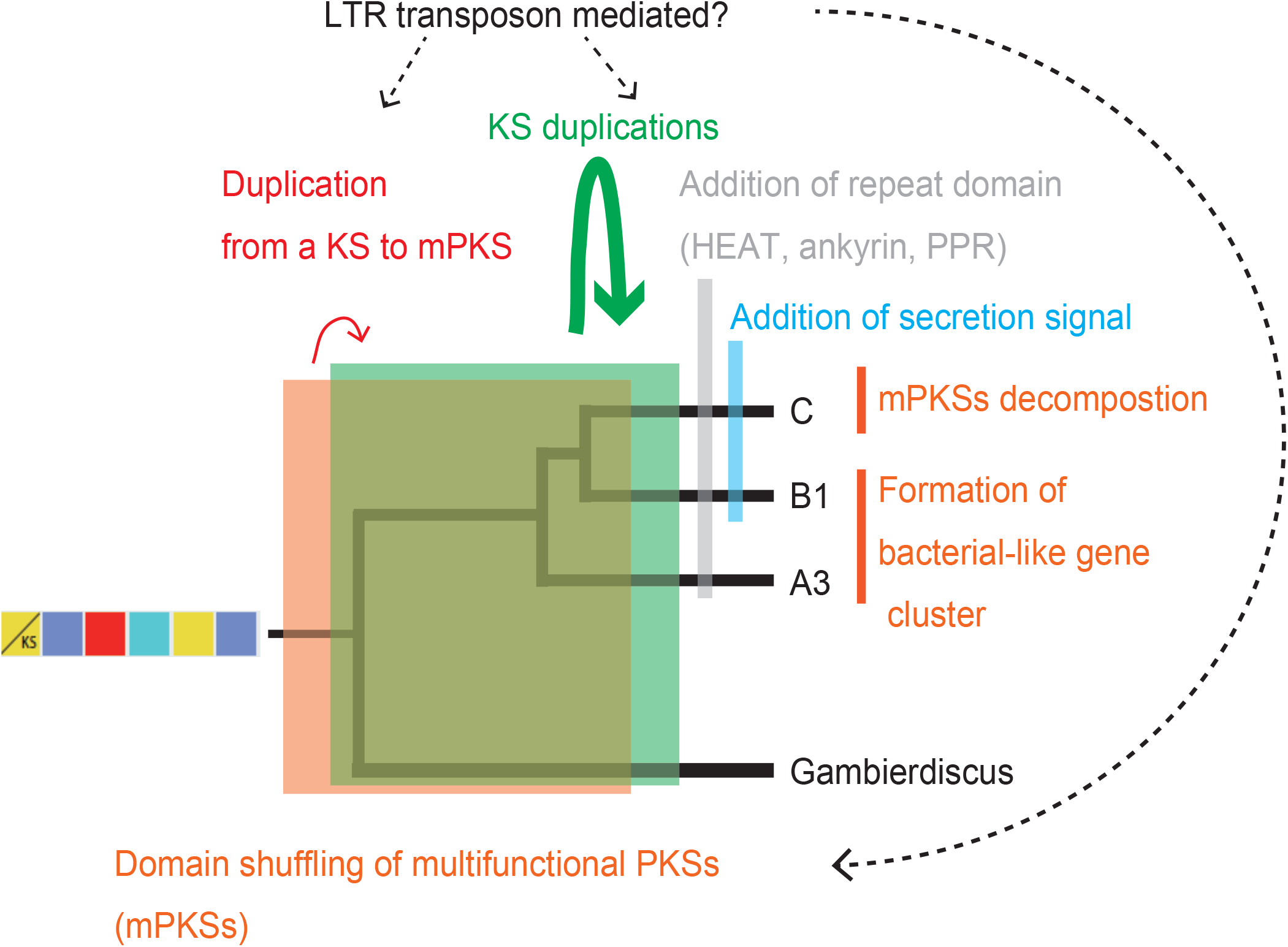
This figure summarizes KS gene evolution in dinoflagellates. Several mechanisms may have contributed to diversification in *Symbiodinium*. Bacterial-like gene clusters can be conserved and retained in several species. Decomposition of such multifunctional polyketide synthases and extensive duplication may have been mediated by LTR transposons, resulting in addition of secretory signals and repeat domains in the three clades (A3, B1 and C).

HGT has been suggested as a significant event contributing to gene innovation with recent evidence linking HGT to various biological processes such as carbohydrate, amino acid, and energy metabolism [59]. HGT is thought to contribute to genome innovation in *Symbiodinium kawagutii*, with 41 out of 56 potential HGT genes being of marine bacterial origin [10]. Gene transfer of PKS genes has been suggested in *Karenia brevis* [60]. Multiple rounds of intra- and intergenic gene duplication have been associated with the expansion of the light-harvesting complex (LHC) gene family in *Symbiodinium minutum* B1, suggesting gene conversion and/or genome rearrangement as an impetus for diversification [61]. Interestingly, monofunctional, probably trans-acting domains of either PKS and NRPS, are often fused with repeats units like HEAT (huntingtin, elongation factor 3, *A* subunit of protein phosphatase 2A and TOR1), ankyrin and pentatricopeptide (PPR) repeats. HEAT repeats have been found in transport-related proteins while the ankyrin repeat family is the second largest dinoflagellate protein family in *Symbiodinium minutum* and is known to facilitate protein-protein interactions involved in several intracellular biological processes [9, 62–64]. On the other hand, PPR proteins are nuclear-encoded, but are targeted to plastids and mitochondria, where they are involved in RNA processing and editing [65–67].

### Evolution of polyketide biosynthesis

Kohli *et al*. [50] suggested that fatty acid synthesis is probably carried out by type II FAS in dinoflagellates, based on a clear distinction between genes involved in fatty acid and polyketide biosynthesis. We found only one gene model (B1030341.t1) associated with type II fatty acid synthesis (FabF-KASII). Our data show that PKS domains have undergone extensive diversification in all three *Symbiodinium* genomes. A plausible explanation for this expansion might be their involvement in novel functions, supported by the fact that ~ 15% of KS and ~9% of AT proteins possess targeting signal peptide, directed towards different organelles. An FAS-like multi-domain polyketide synthase has been identified in *Durinskia baltica* [68], associated with fatty acid biosynthesis. A recent transcriptomic survey of the dinoflagellate *Hematodinium* sp. revealed only type I FAS [69], while another study on *Gambierdiscus* spp. revealed a distinct type II FAS system along with single KS domains [31], suggesting a uniqueness of these pathways to specific dinoflagellates. Both type I and type II FAS systems can exist, as in *Toxoplasma* [70]. Some taxa possess only cytosolic type I, as in *Cryptosporidinium parvum*, while others have only plastid type II, as in *Plasmodium falciparum* [71]. Clearly, apicomplexan and dinoflagellate ancestors possessed both systems.

AT domains of *cis*-AT PKSs display specificity towards various extender units (e.g. methylmalonyl-CoA, hydroxymalonyl-ACP, methoxymalonyl-ACP, etc) while *trans*-ATs are specific for malonyl-CoA. Stand-alone AT proteins have been reported in several PKSs with modules lacking AT domains and these proteins provide malonyl building blocks for the ACP domains of PKS [72, 73]. Our analysis shows that these stand-alone *trans*-AT proteins are dominant in *Symbiodinium* genomes, forming a major clade that may undergo independent evolution compared to canonical *cis*-AT domains. The existence of such *cis-* and *trans*-AT clades has been reported in bacteria and interpreted as proof of independent evolution [74]. Bacterial cis-AT PKS have evolved mainly via module duplication and horizontal/vertical acquisition of entire assembly lines [17], while *trans*-AT has a tendency to recombine and to form novel gene clusters in a mosaic-like fashion [75], as seen in the global pattern of AT in *Symbiodinium* genomes (Fig. 2). Shelest *et al*. [21] found that noniterative PKSs in algae depend mainly on *trans*-AT and are features of multimodular PKS.

### Evolution of non-ribosomal peptide biosynthesis

Few studies have reported NRPS in dinoflagellate transcriptomes [76, 77]; however, detailed analyses of NRPS remain limited. To our knowledge, this is the first study that look at the role and affinities of adenylation and condensation domains in dinoflagellates. Compared to type I PKS, NRPSs were reduced in number within the three *Symbiodinium* clades. NRPSs are less abundant in eukaryotic microalgae [21]. A sequence of amino acids within the A domain catalytic pocket appears to govern recognition and activation of an amino acid substrate. Thus, any point mutations within this segment can drastically change the specificity of the A domain. A mono-modular adenylation domain favors incorporation of polar and non-polar amino acids during peptide synthesis in *Symbiodinium* (Fig. 4a). A conserved domain organization in mono/bi-modular NRPSs exists in fungal species, implying that this architecture is critical for its function [78]. Single A or A-T domains can interact with other NRPS components to achieve biosynthesis by successful activation and transfer of the substrate to the C domain in either the same or different NRPS [79]. NRPSs are mainly modular enzymes with several domains; however, there are reports of nonmodular enzymes among fungal subfamilies [78]. Freestanding A, C, or PCP proteins act *in trans* to form NRPS modules and may be involved in natural product biosynthesis, devoid of the peptide moiety [80].

### Conserved secondary metabolic pathways in *Symbiodinium*

*Symbiodinium* lineages diversified from the ancestral clade A ~50 MYA, at the beginning of the Eocene [81] and have adapted to different environments, performing critical functions in reef ecosystems, as well as serving as photosynthetic endosymbionts of different phyla [7]. New *Symbiodinium* genomes now allow us to compare biosynthetic pathways, shedding light on the organization and role of pathways and their contribution to ecological success. Several gene clusters are conserved between clades A3, B1, and C (Fig. 3b-d), despite the divergence of clades B and C 25-50 and 15 MYA, respectively [81]. Rosic *et al*. [82] reported the importance of conserved phosphatidylinositol signaling pathways in four *Symbiodinium* clades and their contribution to symbiotic interactions. We found that clades A3 and B1 produce unique polyketides, supporting the clade-metabolite hypothesis [5]. Metabolite profiles of different *Symbiodinium* species are influenced by different temperatures and light regimes [82]. On the other hand, metabolomic similarity was detected only between clades B1 and C. At this stage, it is difficult to link specific metabolites to specific pathways, but these results suggest that novel pathways must have evolved in the common ancestor of clades B1 and C so as to provide a common set of metabolites, irrespective of their hosts and environments. Biological systems regulate biochemical and cellular processes when subjected to environmental changes [84]. This study shows that *Symbiodinium* genomes encode PKS and NRPS enzymes with broad substrate tolerance as a cost-effective way of generating chemical diversity. The Screening hypothesis suggest that organisms that produce many chemicals, have more chances of enhanced fitness because greater chemical diversity increases the chance of producing metabolites with unique traits, as illustrated by zooxanthellatoxins and zooxanthellamides [85]. So why are only a few major pathways conserved among these clades? It might be beneficial for organisms to elongate existing pathways to generate new chemical diversity, instead of originating entirely new pathways [86].

## Conclusions

We surveyed three *Symbiodinium* genomes for genes associated with secondary metabolism. We show that PKS genes are more diversified than NRPS genes and that several evolutionary processes have contributed to this diversification. Furthermore, these genes display a degree of substrate specificity and flexibility that has been maintained evolutionarily, irrespective of host system. These results demonstrate that *Symbiodinium* genomes are well equipped to generate chemical diversity when it comes to secondary metabolite biosynthesis. Despite limitations of comparative genomics approaches to assess ecological roles of metabolic pathways, this study provides preliminary insights into how dinoflagellate genomes adapt to host environments and addresses the functional roles of secondary metabolites in such symbiotic relationships.

## Acknowledgements

GB is supported by a Japanese Society for the Promotion of Science (JSPS) Research Fellowship for Young Scientists (DC1). This work was supported partly by JSPS (no. K07454 to E.S) and by generous funding by Okinawa Institute of Science and Technology Graduate University to the Marine Genomics Unit. We thank Steven D. Aird for editing the manuscript. The authors are grateful to Dr. Mary Alice Coffroth and Dr. Michio Hidaka for providing the *Symbiodinium* samples. The authors are thankful to Dr. Chuya Shinzato (The University of Tokyo, Japan) for helpful comments on genomic analysis and to the OIST sequencing and imaging sections for their support.

